# Learning neural decoders without labels using multiple data streams

**DOI:** 10.1101/2021.09.10.459775

**Authors:** Steven M. Peterson, Rajesh P. N. Rao, Bingni W. Brunton

**Affiliations:** Department of Biology, University of Washington, Seattle, WA 98195, USA; eScience Institute, University of Washington; Paul G. Allen School of Computer Science and Engineering, University of Washington; Department of Electrical and Computer Engineering, University of Washington; Center for Neurotechnology, University of Washington

**Keywords:** neural decoding, self-supervised learning, cross-modal learning, deep clustering, electroencephalography

## Abstract

Recent advances in neural decoding have accelerated the development of brain-computer interfaces aimed at assisting users with everyday tasks such as speaking, walking, and manipulating objects. However, current approaches for training neural decoders commonly require large quantities of labeled data, which can be laborious or infeasible to obtain in real-world settings. One intriguing alternative uses self-supervised models that share self-generated pseudo-labels between two data streams; such models have shown exceptional performance on unlabeled audio and video data, but it remains unclear how well they extend to neural decoding. Here, we learn neural decoders without labels by leveraging multiple simultaneously recorded data streams, including neural, kinematic, and physiological signals. Specifically, we apply cross-modal, self-supervised deep clustering to decode movements from brain recordings; these decoders are compared to supervised and unimodal, self-supervised models. We find that sharing pseudo-labels between two data streams during training substantially increases decoding performance compared to unimodal, self-supervised models, with accuracies approaching those of supervised decoders trained on labeled data. Next, we develop decoders trained on three modalities that match or slightly exceed the performance of supervised models, achieving state-of-the-art neural decoding accuracy. Cross-modal decoding is a flexible, promising approach for robust, adaptive neural decoding in real-world applications without any labels.

## 1 Introduction

Brain–computer interfaces that decode neural activity to control robotic or virtual devices have shown great potential to assist patients with neurological disabilities [1–7], while also furthering our understanding of brain function [8–10]. Much of the recent progress in brain-computer interfaces has been driven by advances in neural decoding algorithms [11–14]. However, these algorithms typically rely on supervised learning and thus require large amounts of labeled training data; even as large quantities of neural data are now routinely recorded, generating annotated datasets by curating “ground truth” labels can be laborious and may involve human error [15,16]. Furthermore, decoding algorithms that are useful in the real world must be able to adapt to new scenarios and non-stationary neural signals with few or no labels [17–20]. One promising approach is to use *self-supervised learning* techniques, which generate pseudo-labels from the data itself and then use those pseudo-labels to train a model iteratively without prior labels (see [21, 22] for comprehensive reviews of self-supervised approaches). Self-supervised neural decoding models would ideally avoid overfitting to irrelevant variations in the training data and achieve performance comparable to supervised decoders. Such robust, self-supervised neural decoders would largely eliminate the need for tedious data annotation [23, 24], expedite analyses of large, complex neural datasets, and lay the groundwork for brain-computer interfaces that can dynamically recalibrate in real-world settings.

Self-supervised learning has been most successfully applied in natural language processing and computer vision, with performance similar to that of top supervised models. State-of-the-art techniques in natural language processing are often self-supervised [27–29], using held-out words from the training set to learn robust language models. In computer vision, popular self-supervised approaches include learning low-dimensional representations that reconstruct the original input (e.g. autoencoders [30]) as well as contrastive approaches that learn when augmented image pairs are similar or different (e.g. GANs and Siamese neural networks [31–34]). However, autoencoders typically minimize the mean squared error between the input and reconstruction, which ignores low amplitude, high frequency activity that may be important for the decoding task [35]. For contrastive approaches, creating dissimilar training pairs can be difficult because there are many ways that an image or signal of interest can differ (e.g. comparing an image of a cat’s face with an image of the cat’s paw, the back of the cat’s head, or a dog’s face); in other words, it may not be clear which differences are useful for training a particular model. Several non-contrastive methods have recently been developed that do not require generating dissimilar training pairs [36–39]. For example, deep clustering generates pseudo-labels based on the structure of the data itself, which are then used to iteratively train the model and update the pseudo-labels for the next training step [38]. Many self-supervised techniques have approached or exceeded supervised model performance [33, 36, 37], demonstrating that labeled data is not always necessary to train a robust model.

Even so, self-supervised neural decoding remains a formidable challenge for multiple reasons. First, neural oscillations recorded with scalp electroencephalography (EEG) or intracranial electrocorticography (ECoG) differ greatly from language and image data [40–44]. Relevant features for neural decoding often lack a clear baseline [20,45], are non-stationary over long time periods [46,47], exponentially decrease in amplitude at higher frequencies [48, 49], and occur in a small fraction of the total recording electrodes [50, 51]. Second, while contrastive techniques have been applied for neural decoding [35,52–55], neural data can be noisy and variable from one example to the next, so creating dissimilar examples is often difficult, even with labeled data [56, 57]. Furthermore, many self-supervised approaches in computer vision augment the input images during training to improve model robustness [39,58,59]. While random crops, rotations, and translations make sense for image data, deciding how to augment neural data is less clear, especially when the behaviorally relevant frequencies are not known [52,60]. For these reasons, it is preferable to use a self-supervised approach for neural decoding that does not rely on contrastive learning and data augmentations.

Sharing information across different data streams provides an intriguing opportunity for self-supervised training of neural decoders. Neuroscience research studies often include multiple data streams recorded simultaneously with the neural recordings, such as muscle activation [64, 65], kinematics of human movements [63,66–68], and various physiological signals [69, 70]. Each data stream has its own variability, but behaviorally relevant activity should be evident in multiple data streams, assuming each one is acquired at a similar timescale and contains information related to the behaviors of interest. Therefore, sharing information across multiple data streams plausibly provides realistic variations for robust training without requiring data augmentation. Indeed, multi-modal and sensor fusion approaches are already commonly applied to improve supervised neural decoding performance [24, 71–73]. For self-supervised learning, Alwassel et al. [26] showed that combining audio and video data streams improved the performance of non-contrastive, deep clustering models, with performance matching those of supervised models. However, it is unclear how well this cross-modal approach extends to self-supervised neural decoding.

In this paper, we show that cross-modal, self-supervised training yields state-of-the-art neural decoders that approach supervised decoding performance despite only training on unlabeled data (Fig. 1). We also extend the approach of Alwassel et al. [26] to any number of data streams and assess the performance of neural decoders trained with this cross-modal, self-supervised approach. We compare cross-modal decoders learned without labels to supervised models and unimodal, self-supervised models. We find that sharing pseudo-labels across multiple data streams substantially improves the performance of self-supervised neural decoding models, with accuracies approaching those of supervised models. When we increase the number of data streams from two to three, cross-modal model performance matches or even slightly exceeds supervised model accuracy. This cross-modal, self-supervised approach provides a compelling alternative to tedious data annotation of neural recordings, enabling scalable analyses of large, complex neural datasets and robust brain-computer interfaces that can readily adapt to new real-world scenarios.

**Figure 1:**
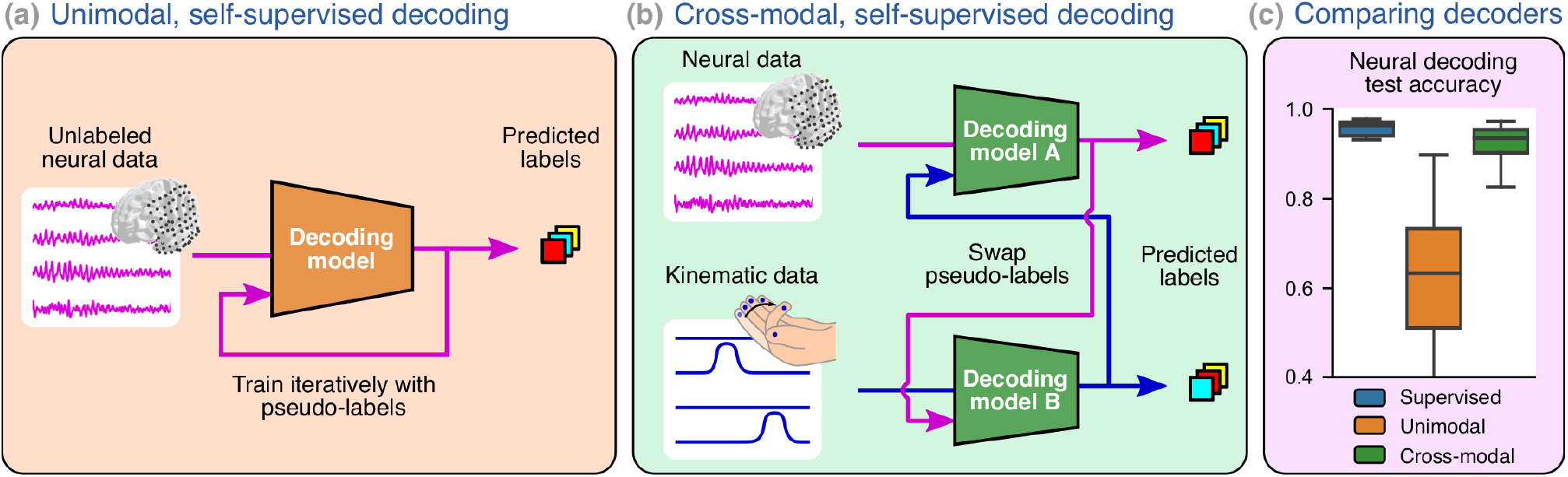
Cross-modal deep clustering for decoding behaviors from unlabeled neural data. **(a)** Self-supervised deep clustering iteratively generates pseudo-labels based on the current decoding model’s outputs in order to train a neural decoder without the need for labeled data. For this study, we use HTNet [25] as the decoding model to predict movement-related behaviors from segmented neural data. **(b)** To improve model robustness, we applied cross-modal deep clustering [26], which swaps neural decoder pseudo-labels with those from additional kinematic and/or physiological data streams. **(c)** We find that neural decoders trained using cross-modal, self-supervised learning approach the performance of supervised neural decoders trained on labeled data.

## 2 Results

Self-supervised learning with deep clustering uses patterns that emerge from the data to iteratively train a neural decoder, and cross-modal deep clustering takes further advantage of correlated patterns among multiple data streams. To understand this training approach, let us first consider a single stream of data, where a unimodal deep clustering decoding model is trained alongside a clustering algorithm that assigns each sample to a cluster based on the decoding model’s output [38, 74]. Training proceeds iteratively, alternating between (1) optimizing the decoding model with the current cluster assignments (pseudo-labels) using backpropagation and (2) updating the pseudo-labels given the current decoding model [74, 75]. Pseudolabels are constrained to equally partition the data to avoid the case where all events are assigned to one cluster. Similarly, cross-modal deep clustering uses pseudolabels, but each decoding model is optimized using pseudo-labels from another data stream. In this way, after many iterations, this swapping of pseudo-labels directly ties together the data streams and the output of their decoding models [26]. Thus, cross-modal deep clustering provides a straightforward procedure to share information among multiple data streams while maintaining separate decoding models for each data stream.

We assessed cross-modal, self-supervised decoding performance on four datasets (Table 1) and demonstrate in each case that cross-modal decoding outperforms unimodal, self-supervised models and approaches the accuracy of supervised models. We consider three movement decoding tasks: determining whether a participant’s arm was moving or at rest (*ECoG move/rest* [56, 76] *and EEG move/rest* [61]), predicting which of five fingers was being flexed (*ECoG finger flexion* [40,62]), and determining whether a participant was exposed to a visual or physical balance perturbation while either walking or standing (*EEG balance perturbations* [63]). Our decoding models all use the HTNet architecture, a compact convolutional neural network that has been demonstrated to perform well at decoding ECoG/EEG data [25, 77].

**Table 1:**
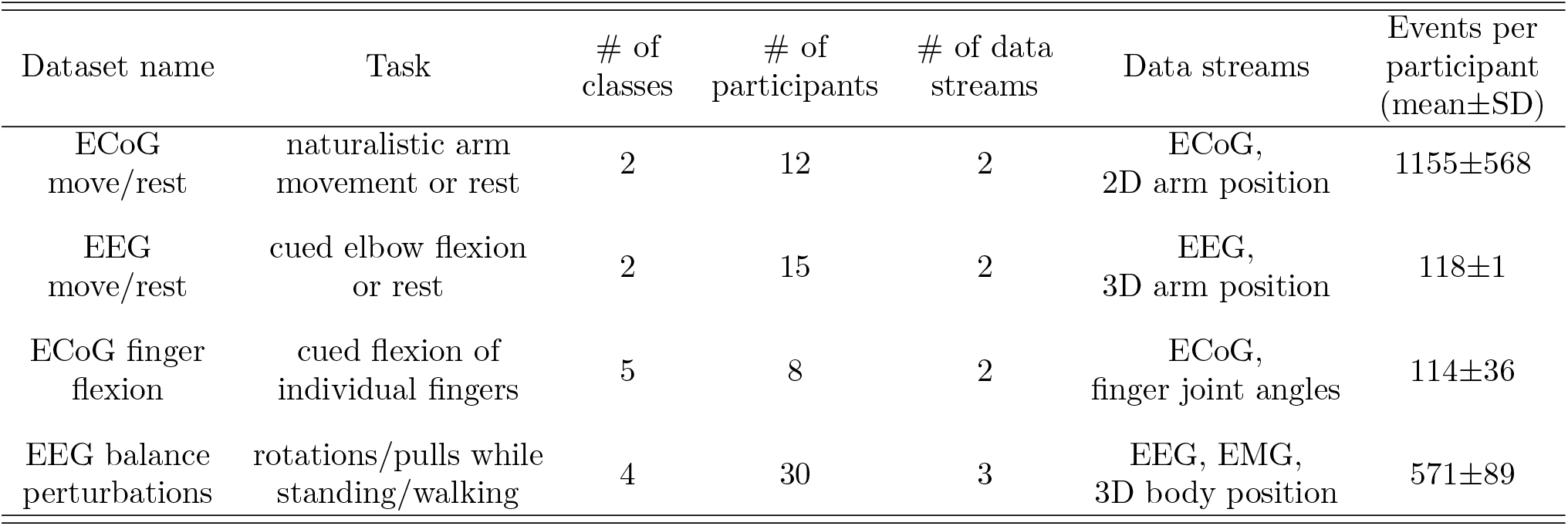
Multi-modal datasets used to test cross-modal neural decoding. We assessed the performance of supervised and self-supervised models using four multi-modal datasets during various movement tasks. Data streams include electrocorticography (ECoG), electroencephalography (EEG), electromyography (EMG), and multiple kinematic measurements. Kinematic measurements were obtained from markerless motion capture applied to video recordings (ECoG move/rest [56]), exoskeleton positions (EEG move/rest [61]), dataglove recordings (ECoG finger flexion [62]), and motion capture markers (EEG balance perturbations [63]). We computed the number of events per participant after balancing events across classes.

| Dataset name | Task | # of classes | # of participants | # of data streams | Data streams | Events per participant (mean±SD) |
| --- | --- | --- | --- | --- | --- | --- |
| ECoG move/rest | naturalistic arm movement or rest | 2 | 12 | 2 | ECoG, 2D arm position | 1155±568 |
| EEG move/rest | cued elbow flexion or rest | 2 | 15 | 2 | EEG, 3D arm position | 118±1 |
| ECoG finger flexion | cued flexion of individual fingers | 5 | 8 | 2 | ECoG, finger joint angles | 114±36 |
| EEG balance perturbations | rotations/pulls while standing/walking | 4 | 30 | 3 | EEG, EMG, 3D body position | 571±89 |

We validated trained decoding models using accuracy on a withheld test set, which reflects how well each model generalizes to unseen data. We also assess each decoder’s ability to effectively cluster unseen data using the v-measure [78]. To compute test accuracy for deep clustering models, we linearly mapped model output clusters to true labels from the training data and used this mapping to generate predictions with the test set [79]. During model training, we found that cross-modal, self-supervised decoders often converged to nearly identical accuracies across data streams for each decoding task (Table S1). The differences in test accuracy that we report in this section primarily reflect how well each trained model is able to generalize to unseen data from its respective data stream.

### 2.1 Cross-modal decoding with two data streams

For all decoding tasks, we find that sharing self-supervised pseudo-labels among two data streams substantially improves decoding performance compared to unimodal, self-supervised models, while also approaching test accuracies of supervised models. In addition, we observe similar differences across model types for clustering performance (Table S2).

#### 2.1.1 Decoding arm movement vs. rest

In both move/rest decoding tasks, cross-modal, self-supervised neural decoders consistently outperform unimodal, self-supervised models and approach supervised model test accuracy (Fig. 2). For the naturalistic ECoG move/rest dataset, we find that neural decoder test accuracy is significantly affected by model type (*p* = 3.25*e* − 5; Friedman test [80]). Cross-modal ECoG decoders achieve an average accuracy of 87% ± 8% (mean±SD), which is well above random chance (50%). We find that cross-modal decoders have a small but significant decrease in test accuracy compared to supervised decoding performance (91%±5%, *p* = 0.005; Wilcoxon signed-rank test with false discovery rate correction [81, 82]). In contrast, the average test accuracy for unimodal, self-supervised decoders is only 57% ± 5%, which is significantly lower than both cross-modal and supervised model performances (*p* = 7.32*e* − 4 for both). We find similar differences among models for the EEG move/rest dataset, with EEG test accuracy significantly affected by model type (*p* = 1.73*e* − 6). Again, the test accuracies of cross-modal (74% ± 10%) and supervised (81% ± 8%) decoders are well above random chance (50%), but do significantly differ from each other (*p* = 0.003). Still, both cross-modal and supervised decoders substantially outperform unimodal, self-supervised models (50% ± 6%, *p* = 9.20*e* − 5 for both comparisons), demonstrating the usefulness of cross-modal training for self-supervised neural decoding.

**Figure 2:**
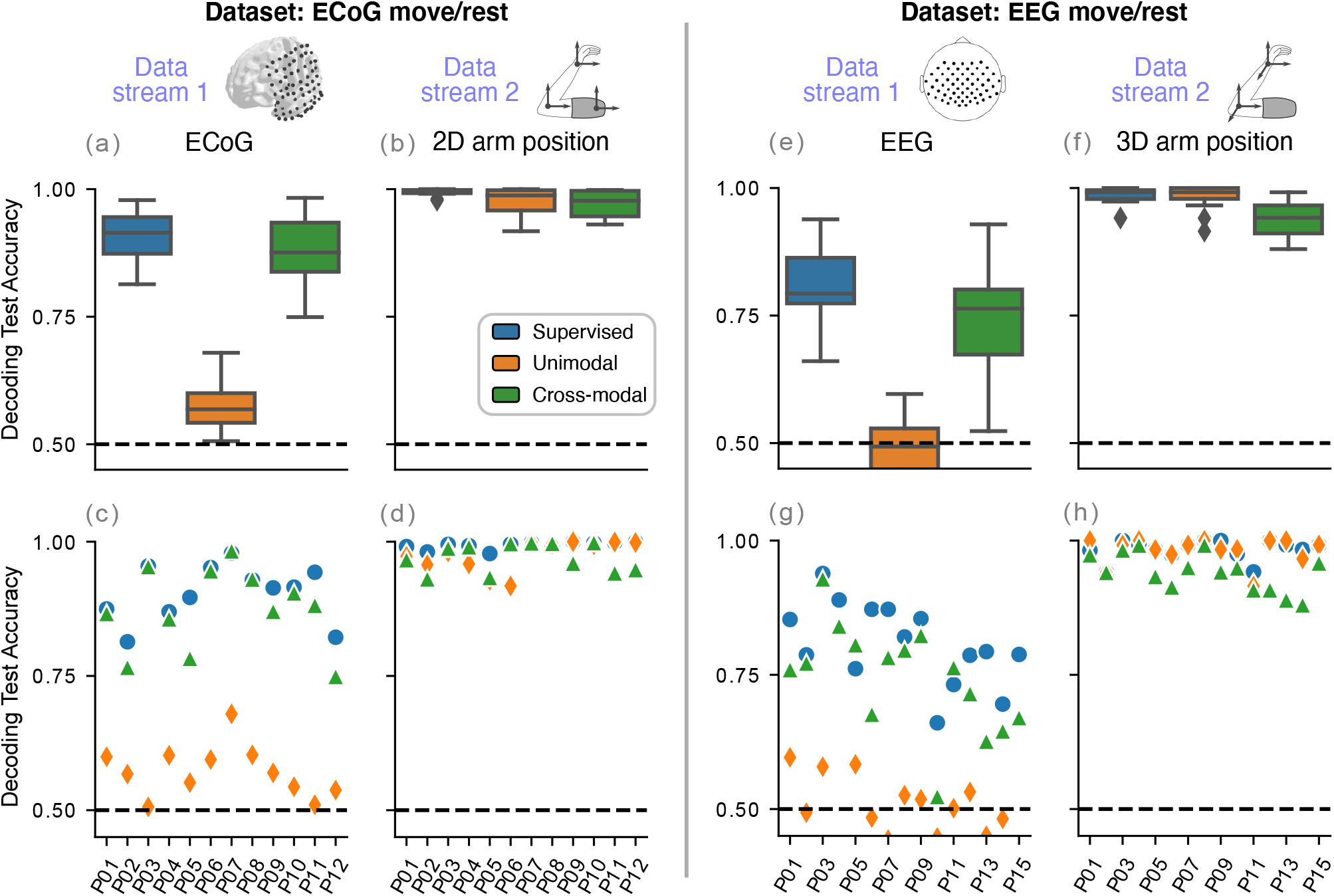
Cross-modal decoding of arm movement vs. rest approaches supervised decoding accuracies. Decoding test accuracy is shown for ECoG move/rest **(a–b)** and EEG move/rest **(e–f)** datasets, averaged over 10 random folds. Note that cross-modal training accuracy is quite similar between the two decoders from each data stream; the differences in test accuracy primarily reflect how well each decoder is able to generalize to unseen data from its respective data stream. For both datasets, the cross-modal decoders are able to leverage the high separability in the arm position data stream to achieve test accuracies well above unimodal, self-supervised models that approach supervised model performance. Single-participant performance is shown in **(c–d)** and **(g–h)** for the ECoG move/rest and EEG move/rest datasets, respectively. Random chance performance is denoted by the dashed, horizontal line at 50%.

For both move/rest tasks, all three model types decode unseen arm position with over 90% accuracy. We find that this pose decoding performance is significantly affected by model type for EEG move/rest (*p* = 9.65*e* − 6), but not for the ECoG move/rest dataset (*p* = 0.067). For ECoG move/rest, supervised models are the most accurate at decoding move/rest from 2D arm position (99% ± 1%), followed by unimodal (98% ± 3%) and cross-modal (97% ± 3%) models. When decoding 3D arm position for the EEG move/rest dataset, we find that cross-modal models (94%±4%) have a small but significant decrease in decoding performance compared to supervised (98%±2%, *p* = 0.002) and unimodal (98% ± 2%, *p* = 0.002) models. These high test accuracies for all decoders indicate how separable the move and rest classes are when decoding arm position. Cross-modal models are able to leverage this high separability in the arm position data stream to improve self-supervised neural decoding performance, which explains why cross-modal neural decoders notably outperform unimodal, self-supervised models.

#### 2.1.2 Decoding finger flexion

When testing decoding performance on a more complex task, we find that pairwise cross-modal training again improves neural decoding performance substantially compared with unimodal, self-supervised models and nears supervised model performance (Fig. 3). We observe a significant effect of model type on neural decoding for this task (*p* = 0.002). Cross-modal neural decoders (53%±12%; mean±SD) achieve substantially higher test accuracies than unimodal, self-supervised models (23%±4%) and approach supervised model performance (57%±9%). Unimodal neural decoders perform near random chance (20%) for most participants, which is significantly worse than both cross-modal and supervised neural decoders (*p* = 0.012 for both).

**Figure 3:**
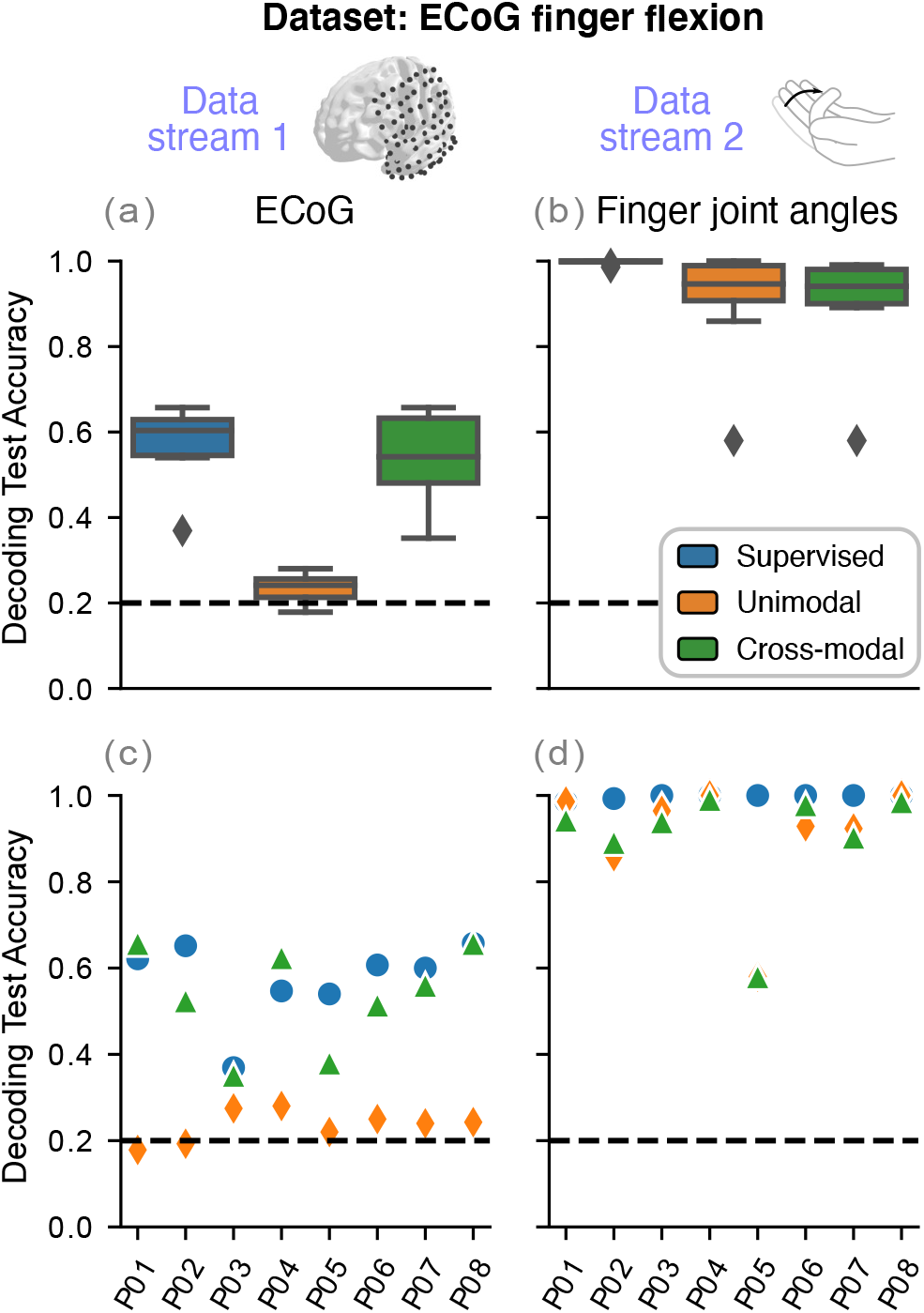
Cross-modal neural decoders achieve comparable performance to supervised models for decoding finger flexion. **(a–b)** Cross-modal decoders again leverage the high separability in finger joint angles for different finger movements to nearly match supervised neural decoding performance. **(c–d)** Decoding performance is shown for each participant, averaged over 10 random folds. Random chance is indicated by the dashed, horizontal line at 20%.

Similar to decoding arm position for the move/rest datasets, all three model types can decode unseen finger joint angles at or above 90% accuracy. We observe a significant effect of model type on test accuracy when decoding joint angles (*p* = 0.008). Cross-modal joint angle decoders (90% ± 14%) have a small but significant decrease in performance compared to supervised models (100%±1%, *p* = 0.023). Unimodal, self-supervised model accuracy (91% ± 14%) is similar to cross-modal performance, indicating that supervised training yields more robust joint angle decoders than self-supervised learning. One explanation is this dataset contains a low number of events per class (23±7 events per class) and that adding more training data would close the performance gap between supervised and self-supervised models. Regardless, cross-modal training is again able to leverage the high separability within the finger joint angle data stream to train high-quality neural decoders.

#### 2.1.3 Decoding balance perturbations

For decoding balance perturbations, our results are consistent in observing that the pairwise cross-modal neural decoders outperform unimodal, self-supervised models and approach supervised model performance (Fig. 4). Because many participants performed this task, almost all comparisons of decoder test accuracy are statistically significant (*p <* 0.05), except between the two pairwise cross-modal models for the EEG data stream (*p* = 0.154). Still, we find that the pairwise cross-modal neural decoders (EEG/body pose: 79% ± 18%; EEG/EMG: 77% ± 19%) perform in between unimodal, self-supervised models (62% ± 18%) and supervised models (88% ± 24%). Note that these average test accuracies are heavily influenced by poor decoding performance near random chance (25%) for P02, P04, P11, and P15. These and other neural decoding outliers reflect the variable signal quality across participants during mobile EEG recordings, especially with only minimal pre-processing to reduce noise. Even so, pairwise cross-modal neural decoders using either body position or EMG clearly outperform unimodal, self-supervised models and demonstrate that any task-relevant data stream can be useful in improving self-supervised decoding performance.

**Figure 4:**
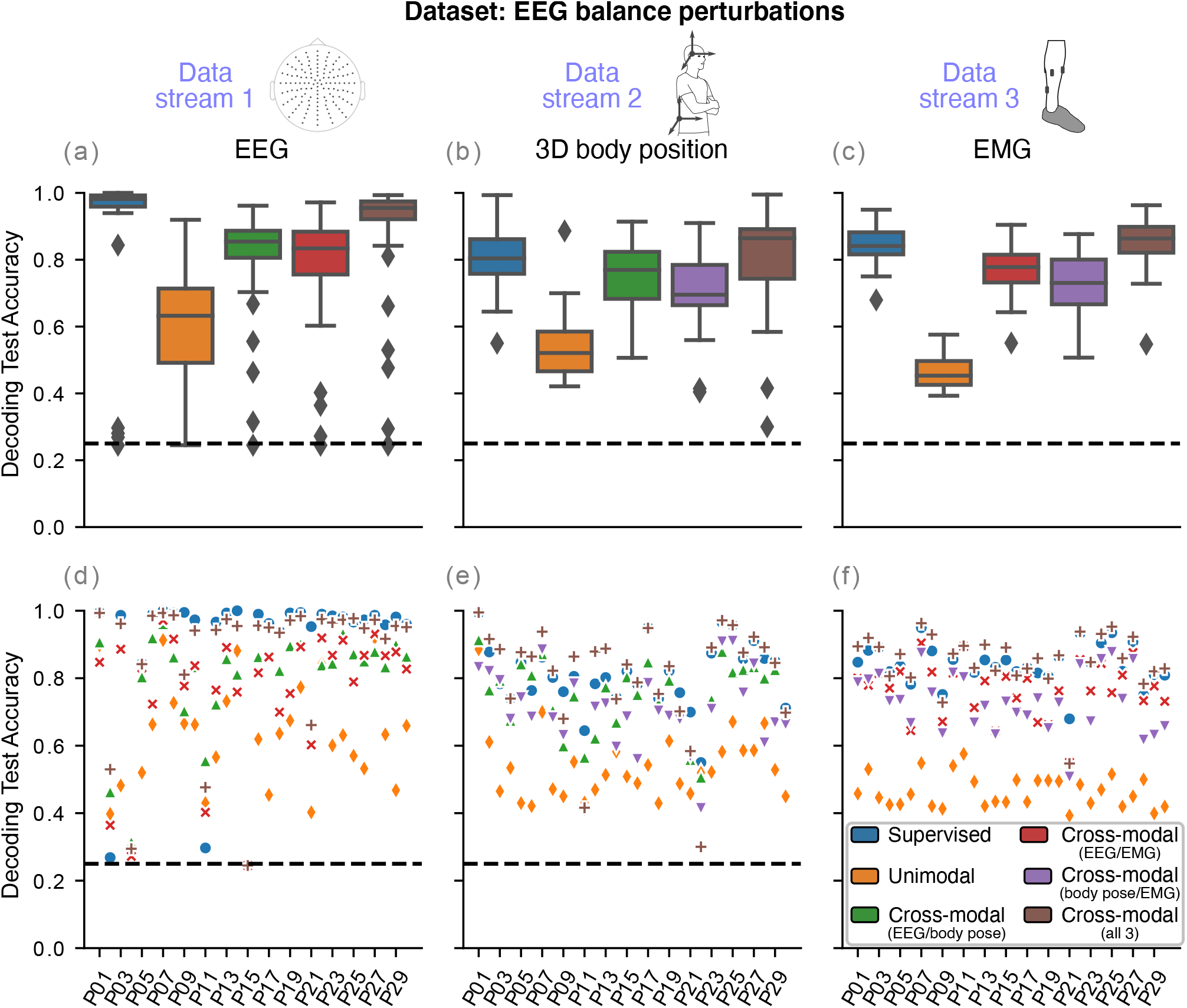
Cross-modal decoding across three data streams matches or even exceeds supervised model performance. **(a–c)** We performed cross-modal decoding using two or all three data streams. Cross-modal models using all three data streams perform similar to or slightly better than supervised decoders, demonstrating that sharing information among multiple data streams can lead to robust neural decoders even without labeled training data. **(d– f)** Single-participant test accuracies show the high variability decoding performance, especially for the EEG data stream. This high variability in EEG decoding performance is not surprising because recordings took place during perturbed balance-beam walking and are likely contaminated with noise from head motion, neck muscles, and eye movements. Random chance is denoted by the dashed, horizontal line at 25%.

Unlike the other datasets, pairwise cross-modal neural decoders approach supervised model performance despite using a non-neural data stream that is not easily separable across classes. Supervised decoding performance is well above chance (body position: 81% ± 10%; EMG: 84% ± 6%), but not at the near-perfect test accuracies seen for the other datasets. Similarly, unimodal, self-supervised models only achieve ∼ 50% test accuracy (body position: 54%±10%; EMG: 46%±5%), which is far below pairwise cross-modal accuracies for decoding body position (EEG/body pose: 75% ± 10%; body pose/EMG: 70% ± 12%) and EMG (EEG/EMG: 77%±8%; body pose/EMG: 73%±9%). These results demonstrate how cross-modal training can still develop high-quality self-supervised decoders, even without an easily separable data stream.

### 2.2 Cross-modal decoding with three data streams

By combining all three data streams from the EEG balance perturbations dataset, we improve cross-modal decoding performance to nearly match or even slightly exceed supervised model performance (Fig. 4). Again, almost all comparisons are statistically significant (*p <* 0.05) due to how many participants are in this dataset. For decoding EEG, trimodal neural decoding performance (87% ± 21%) is substantially higher than unimodal and pairwise cross-modal accuracies. We do find a small but significant decrease in trimodal performance compared with supervised decoders (*p* = 0.003), but both decoders are within ∼ 1–3% for most participants. In addition, trimodal decoding performance for the non-neural data streams is notably increased compared to pairwise cross-modal decoders (body position: 81% ± 16%, EMG: 85% ± 8%). For decoding body position, trimodal decoding performance is not significantly different from supervised decoder accuracies (*p* = 0.133). For EMG decoding, we actually observe a small but significant increase in trimodal decoding accuracy over supervised decoding performance (*p* = 0.005). We also find similar differences among model types for clustering performance (Table S3). Taken together, our findings demonstrate that including additional data streams can lead to robust self-supervised decoders that do not require labeled training data.

### 2.3 Effect of expected number of clusters on performance

We also assess how the expected number of clusters (K) impacts pairwise cross-modal performance, finding that selecting a value for K that is above the true number of classes notably affects test accuracy but not clustering performance (Fig. S1). We performed this assessment on the ECoG finger flexion dataset. For both data streams, setting K to less than the number of classes led to decreases in test accuracy and clustering performance relative to cross-modal models with K equal to the number of classes. In addition, we found that over-clustering, or setting K to greater than the number of classes, also decreases test accuracy compared with K equal to the number of classes, likely because the model had difficulty generalizing clusters that divided up the same class. In contrast, clustering performance for over-clustered models increases or stays the same compared to models with K equal to the number of classes. These findings highlight the importance of carefully selecting K depending on the overall goal; if generalized decoding is desired, then K should be carefully chosen, but if clustering is the primary objective, then choosing a large K should be sufficient.

## 3 Discussion

In this paper, we demonstrate that cross-modal deep clustering extends well to neural decoding applications. We show four examples where cross-modal neural decoders achieve performance that approaches supervised models despite using no labeled data. Because neural recordings are routinely measured simultaneously with multiple other data streams, we also extend this cross-modal approach to include more than two data streams and find notably improved performance when adding a third data stream to train cross-modal neural decoders. Our findings demonstrate that including additional, task-relevant data streams during cross-modal training leads to robust, self-supervised neural decoders that do not require labeled data.

To our knowledge, cross-modal deep clustering is the first self-supervised approach that can create high-performing neural decoders from unlabeled data without relying on contrastive learning and data augmentations. This approach seems to perform well because variations among different, task-relevant data streams reduce overfitting during model training, similar to what is achieved by data augmentation [52, 53]. Even supervised neural decoders can overfit to the inherent variability of neural data, as well as task-irrelevant changes in neural activity due to factors such as fatigue and emotional state [14, 83, 84]. These task-irrelevant patterns are less likely to appear across multiple data streams, thus reducing the tendency to overfit. This hypothesis also explains why adding more data streams improves cross-modal decoder performance for the EEG balance perturbations dataset (Fig. 4) and underlies many sensor fusion and multi-modal approaches for reliable neural decoding [71–73, 85, 86]. Our proposed approach for cross-modal deep clustering using any number of data streams (Fig. 5) leverages the multiple neural, physiological, and kinematic data streams often collected in neuroscience research studies [24, 66–70, 87, 88].

**Figure 5:**
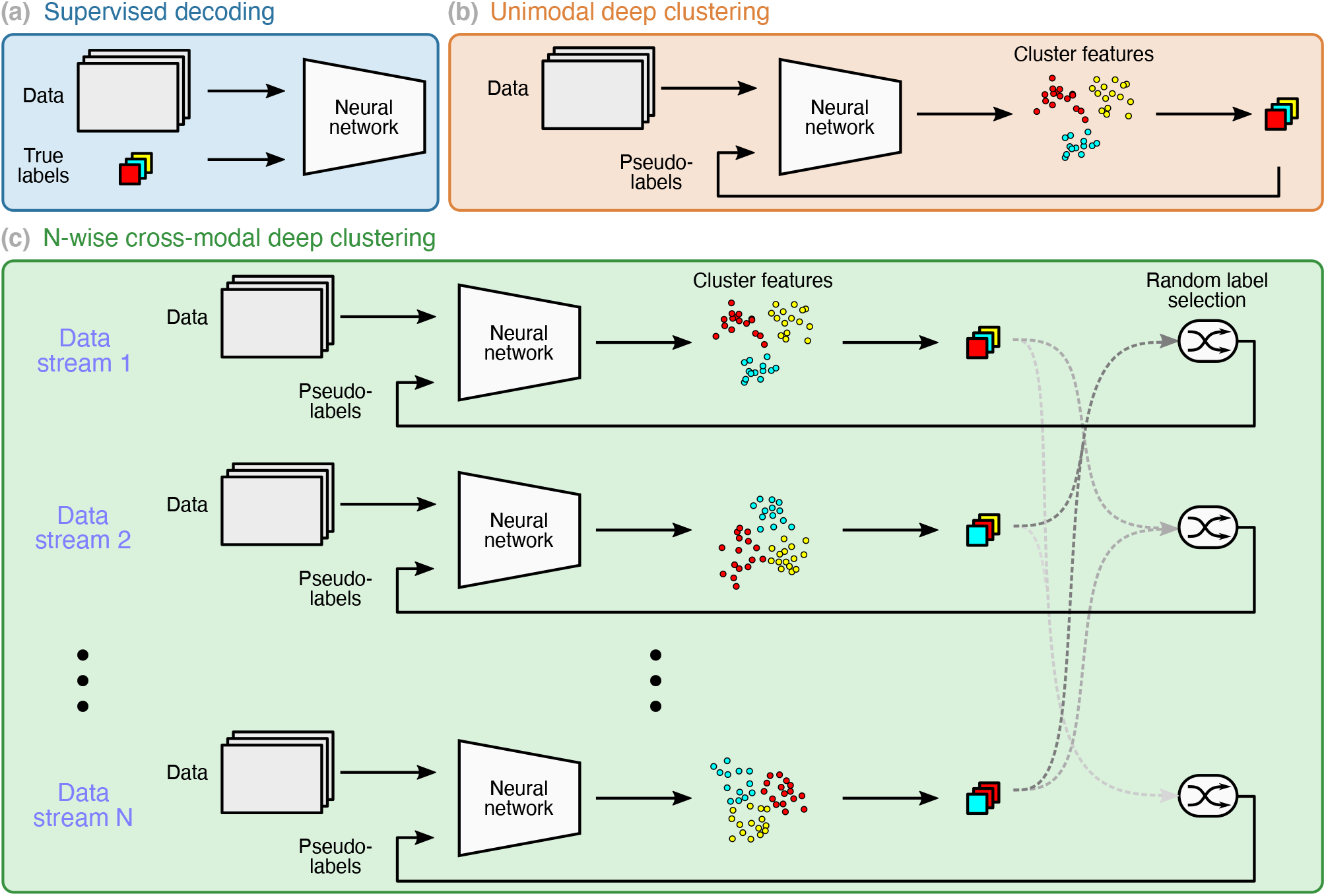
Supervised and self-supervised training paradigms. We compare performance of supervised neural decoders with unimodal and cross-modal self-supervised deep clustering models. **(a)** Supervised decoders are trained using the true data labels. **(b)** In contrast, deep clustering self-generates pseudo-labels by clustering the neural network model outputs and then uses these pseudo-labels to train the model. Note that the neural network and clustering technique are alternately optimized during training. **(c)** In cross-modal deep clustering, decoder models for each data steam are trained using pseudo-labels randomly selected from all other data streams except itself. For all supervised and self-supervised approaches, we used HTNet, a convolutional neural network for decoding ECoG and EEG data, as the neural network decoder.

Cross-modal deep clustering can also be adapted to a wide variety of different model architectures and data streams. Here, we used a convolutional neural network (HTNet) for decoding [25], but we could have applied a larger convolutional neural network model or added recurrent layers [89, 90]. Such adjustments could potentially improve decoding accuracy, but would increase training time and possibly require more training data to train. HTNet is also able to pool neural data across multiple participants, raising the possibility of training neural decoders on unlabeled, multi-participant data [25]. In addition to its flexibility in model architecture, cross-modal deep clustering can also be applied to any data streams that are synchronously recorded. We chose kinematic data streams such as body position and joint angle because of their high relevance to movement behaviors [91]. Ideal data streams contain high-quality, task-relevant information. For exploratory analyses of large neural datasets, selecting certain data streams could also be used to bias models towards decoding specific behaviors.

Despite its impressive decoding performance on unlabeled data, cross-modal deep clustering has multiple limitations to consider. First, like with many clustering algorithms, the number of expected clusters must be chosen with extreme care to obtain high-quality decoding performance on unseen data (Fig. S1). One way to decide on a reasonable number of expected clusters without data labels is to generate clusters for a subset of examples, train a classifier using the cluster assignments, and test how well this classifier’s predictions match the cluster assignments for the unseen data. This cross-validation approach has been used for selecting the optimal number of clusters during k-means clustering [92]. Another limiting factor of our current approach is that it requires segmented, event-related neural data that transitions from a consistent baseline rest state. We could have potentially extended our approach to continuous data using a sliding window [93, 94], but we would have to account for event transitions from non-rest states [95]. Finally, our approach may not be appropriate for decoding imagined movements, where kinematic data streams are not available. Still, studies have shown that other data streams such as eye gaze, heart rate, and muscle activity can be affected by imagined movements [96–98] and thus may be useful for cross-modal decoding of imagined movements.

This cross-modal deep clustering approach can be extended to a wide variety of neural decoding applications where limited labeled data is available, including predicting mood and somatosensation [99, 100]. In addition, fine-tuning and other transfer learning techniques can be readily applied to improve trained model performance, as is often done for self-supervised models [52, 53]. Our current approach of randomly sharing pseudo-labels for more than two data streams may be made more rigorous by weighting each data stream based on its reliability and data quality. Data augmentations could also be useful in improving decoder performance. While we focused on self-supervised learning without data augmentations, deep clustering has been frequently applied to augmented data [38, 39, 74]. Using multiple data streams should help minimize any biases introduced by a specific data augmentation. Overall, cross-modal deep clustering is a promising alternative to tedious data annotation and provides a reliable framework for achieving high-quality neural decoding when curating an annotated dataset is difficult or infeasible.

## 4 Methods

### 4.1 Cross-modal deep clustering

We implemented the unimodal deep clustering approach from Asano et al. [74] (Fig. 5(b)). This unimodal approach alternates between (1) optimizing the decoding model given the current pseudo-labels and (2) updating the pseudo-labels given the current decoding model. This first step involves minimizing cross-entropy loss, similar to supervised training (Fig. 5(a)). In the second step, the Sinkhorn-Knopp algorithm takes in the model outputs and updates the label assignments with the constraint that the pseudo-labels must equally partition the data among the expected number of clusters [74, 75]. This constraint avoids situations where all events are assigned the same pseudo-label and the model minimizes cross-entropy loss by always predicting the same label [74]. We primarily set the number of expected clusters (K) equal to the number of true clusters and later explored the effects of K on model performance. All models were created and trained using Python 3.8.5 and PyTorch 1.7.1.

We modified this unimodal approach from Asano et al. [74] to share pseudo-labels across multiple modalities as done in Alwassel et al. [26] (Fig. 5(c)). This cross-modal deep clustering approach was found to out-perform other multi-modal deep clustering models [26]. Cross-modal deep clustering involves swapping pseudo-labels between the two data streams when optimizing the decoding model. This swapping directly affects not only how the decoding model is learned, but also indirectly what the pseudo-labels are because they are generated from the decoding model’s output.

We extend this pairwise cross-modal deep clustering approach to accommodate more than two data streams. For each event, we randomly select a pseudo-label to train the decoding model for a particular data stream from any of the other data streams. When there are only two data streams, the two models simply swap pseudo-labels, just like the previous pairwise approach [26]. With our extended N-wise cross-modal deep clustering approach, we assume that the decoding models for every data stream will converge to similar pseudo-labels by the end of training. Otherwise, the trained models may be biased to the most recent pseudo-labels used during training.

### 4.2 HTNet decoding model

For the decoding model, we used HTNet [25], a convolutional neural network for generalized neural decoding. We chose HTNet because it has few parameters and identifies data-driven spatiotemporal features in the frequency domain, where neural data is often most easily separated. We also applied HTNet for decoding all non-neural data streams to maintain consistency across datasets. We used the user-defined number of expected clusters to determine the size of HTNet’s output layer. That way, HTNet generates one value for each expected cluster, which are transformed using the softmax function into joint probabilities estimated by the model. These estimated probabilities are compared to the pseudo-label for event and cross-entropy loss between the two is minimized [74].

We slightly modified the original HTNet architecture by replacing the initial temporal convolution layer with SincNet filters [101]. Instead of learning the entire convolution kernel, each SincNet filter learns only the minimum and maximum frequency cutoffs for a pre-defined band-pass filter. Compared with a temporal convolution layer, SincNet filters are less biased towards low frequencies because they do not need to learn the complex kernel shape often needed for isolating high frequencies. We found that SincNet filters improve HTNet’s performance, especially for data streams such as ECoG with task-relevant features at high-frequencies.

### 4.3 Datasets

We assess cross-modal deep clustering performance on four datasets that span a variety of tasks and recording modalities (Table 1). All four datasets are publicly available and include at least one other synchronized data stream besides the neural recording. Note that we did not discard noisy events for any dataset. Variability in the number of events across participants reflects either differences in the number of events actually recorded or events that were discarded because not enough data was available to segment.

#### 4.3.1 ECoG move vs. rest dataset

Concurrent intracranial ECoG and upper-body positions were obtained from 12 human participants (8 males, 4 females; aged 29±8 years old [mean±SD]) during continuous clinical epilepsy monitoring conducted at Harborview Medical Center in Seattle, WA [56, 76, 102]. ECoG electrodes were implanted primarily in one hemisphere (5 right, 7 left) and included cortical surface recordings along with penetrating electrodes into subcortical areas. This study was approved by the University of Washington Institutional Review Board, and all participants provided written informed consent. ECoG and pose data were sampled at 500 Hz and 30 FPS, respectively. This dataset is publicly available at: https://doi.org/10.6084/m9.figshare.16599782.

The goal is to discriminate between rest events and movements of the arm contralateral to the implanted electrode hemisphere. Move/rest classifications were determined via markerless pose tracking of video recordings and automated state segmentation [76]. Move events were defined as wrist movements that followed 0.5 seconds of no movement. In contrast, rest events denoted periods of 3 or more seconds with no movement in either wrist. We balanced the number of move/rest events using random under-sampling [103], leading to 1155±568 events per participant.

ECoG data was preprocessed by removing DC drift, high-amplitude discontinuities, band-pass filtering (1– 200 Hz), notch filtering, referencing to the common median across electrodes, and downsampling to 250 Hz [56]. We selected two-dimensional pose trajectories obtained from video recordings for 3 keypoints (shoulder, elbow, and wrist of the contralateral arm) and computed the difference between neighboring time-points. Data from both modalities were trimmed to 2-second segments centered around each event.

#### 4.3.2 EEG move vs. rest dataset

Concurrent EEG and arm positions were recorded from 15 human participants during visually cued elbow flexion (6 males, 9 females; aged 27±5 years old [mean±SD]). EEG data was recorded from 61 electrodes and sampled at 512 Hz. For pose data, we selected three-dimensional elbow and wrist positions recorded by an exoskeleton during the experiment and sampled at 512 Hz. This study was approved by the ethics committee of the Medical University of Graz, and all participants provided written informed consent [61]. This dataset is publicly available at: http://bnci-horizon-2020.eu/database/data-sets.

Similar to the previous dataset, the decoding task involves 2-class classification of either move or rest events. Here, a move event corresponds to cued elbow flexion of the right arm. Each participant performed 120 total trials (60 movement and 60 rest trials). We aligned data segments to movement onset, which was identified by thresholding the wrist’s radial displacement after the visual cue to move.

EEG data was notch filtered at 50 Hz, referenced to the right mastoid, and band-pass filtered (0.01–200 Hz). We performed further processing by high-pass filtering at 1 Hz, average referencing, and resampling to 250 Hz. Three-dimensional pose trajectories were obtained from each participant’s right elbow and hand, and we computed the difference between neighboring timepoints to quantify the change in position.

#### 4.3.3 ECoG finger flexion dataset

ECoG and finger joint angles were recorded concurrently from 9 human participants during visually cued finger flexion (3 males, 6 females; aged 27±9 years old). ECoG electrodes were implanted primarily in one hemisphere (2 right, 7 left) and included only cortical surface recordings (38–64 electrodes per participant). Finger joint angles were recorded using a 5 degree-of-freedom dataglove sensor [62]. All patients participated in a purely voluntary manner, after providing informed written consent, under experimental protocols approved by the Institutional Review Board of the University of Washington (#12193). All patient data was anonymized according to IRB protocol, in accordance with HIPAA mandate. These data originally appeared in the manuscript “Human Motor Cortical Activity Is Selectively Phase-Entrained on Underlying Rhythms” published in PLoS Computational Biology in 2012 [62]. Both ECoG and finger joint angles were originally sampled at 1000 Hz. This dataset is publicly available at: https://searchworks.stanford.edu/view/zk881ps0522 [40].

The decoding task here is to classify which of five fingers is being flexed. All finger flexions occurred in the hand contralateral to the ECoG implantation hemisphere. Each cued finger flexion lasted 2 seconds, followed by 2 seconds of rest. Every participant performed 150 pseudo-randomly interleaved finger flexions (30 for each finger).

We discarded one participant due to an inability to obtain cued movement times. ECoG data was band-pass filtered between 4–250 Hz, notch filtered at 60 Hz and its harmonics, average referenced, and downsampled to 250 Hz. Pose data was standardized and also downsampled to 250 Hz. We removed finger flexion events where no movement occurred.

#### 4.3.4 EEG balance perturbations dataset

Concurrent EEG, electromyography (EMG), and three-dimensional body position were collected from 30 human participants during sensorimotor perturbations to standing and walking balance (15 males, 15 females; aged 23±5 years old [mean±SD]) [63]. EEG data was recorded from 128 electrodes and sampled at 512 Hz. EMG was sampled at 1 kHz and collected from 4 muscles on each leg (8 total): tibialis anterior, soleus, medial gastrocnemius, and peroneus longus. Three-dimensional body position of the head and sacrum was recorded using motion capture markers with 100 Hz sampling. This protocol was approved by the University of Michigan Health Sciences and Behavioral Sciences Institutional Review Board, and all subjects provided written informed consent [63]. This dataset is publicly available at: https://openneuro.org/datasets/ds003739.

The decoding task for this dataset is to classify between two sensorimotor perturbations during either standing or walking (4 total classes). Sensorimotor perturbations involved either a 1-second mediolateral pull at the waist or a half-second 20 degree field-of-view rotation using a virtual reality headset. Each class includes 150 events recorded during a separate 10-minute recording session and balanced between left/right pulls or clockwise/counterclockwise rotations.

EEG data were downsampled to 256 Hz, 1 Hz high-pass filtered, and referenced to the common median across electrodes. We removed the linear trends in EMG and pose data using Matlab 2013a. EMG and pose data were then resampled to match the EEG sampling rate of 256 Hz.

### 4.4 Model comparisons and hyperparameters

We compared decoder model performance after three types of training: supervised, unimodal self-supervised, and cross-modal self-supervised. All decoder hyperparameters were fixed across these three models; the only difference was using either the true labels or self-generated pseudo-labels during model training. We trained supervised and unimodal, self-supervised models for 40 epochs, while cross-modal models were trained for 200 epochs. We selected 40 epochs based on the average number of epochs used to train HTNet on similar neural data [25]. For that approach, the likelihood of overfitting was minimized by applying early stopping based on improvement in validation set accuracy. For cross-modal models, we performed model training over many more epochs in order to provide enough time to converge across all data streams. Unlike unimodal and supervised models, we found that cross-modal models did not overfit when trained over hundreds of epochs.

We set decoder model (HTNet) hyperparameters for the two move/rest datasets based on our previous decoding study [25]. For the ECoG finger flexion encoder, we increased the number of spatial filters from 2 to 5 (Table S4) to adequately capture different spatial features across finger movements. To keep the total number of fitted parameters low, we decreased the number of temporal filters to 6. In addition, we didn’t use SincNet and spectral power for the arm position data to preserve the low-frequency features of the data. For the EEG balance dataset, we similarly increased the number of spatial filters to 3 and decreased the number of temporal filters to 6.

### 4.5 Model validation metrics

We chose two metrics for validating trained decoding model performance: test accuracy and v-measure. Both model validation metrics are always averaged over 10 folds, selected with stratified random sampling to preserve the percentage of samples for each class within each folds. Test accuracy measures model classification accuracy on unseen data, while v-measure assesses the model’s ability to separately cluster different classes. For accuracy, we computed the linear mapping between true and predicted clusters that yielded the maximum accuracy on the training set [79]. Then, we computed the accuracy on the test set using the same mapping between true and predicted clusters. For v-measure, we computed the harmonic mean of completeness (how much of each class is present in a single cluster) and homogeneity (how much each cluster is made up of a single class) [78]. V-measure can range between 0 and 1, with 1 indicating perfect clustering. We focus primarily on test accuracy instead of clustering performance because we want trained decoding models that can generalize to unseen data.

## Code and data availability

Our code is publicly available at: https://github.com/BruntonUWBio/cross-modal-ssl-htnet. The code in this repository can be used in conjunction with publicly available ECoG [40, 76] and EEG [61, 63] datasets to generate all of the main findings and figures from our study.

## Acknowledgements

We thank Satpreet Singh, Zoe Steine-Hanson, Ellie Strandquist, and Pierre Karashchuk for discussions and help with data analysis and model design. This research was supported by funding from the National Science Foundation (1630178 and EEC-1028725), the Washington Research Foundation, and the Weill Neurohub.

## 6 Supplementary Information

**Figure S1:**
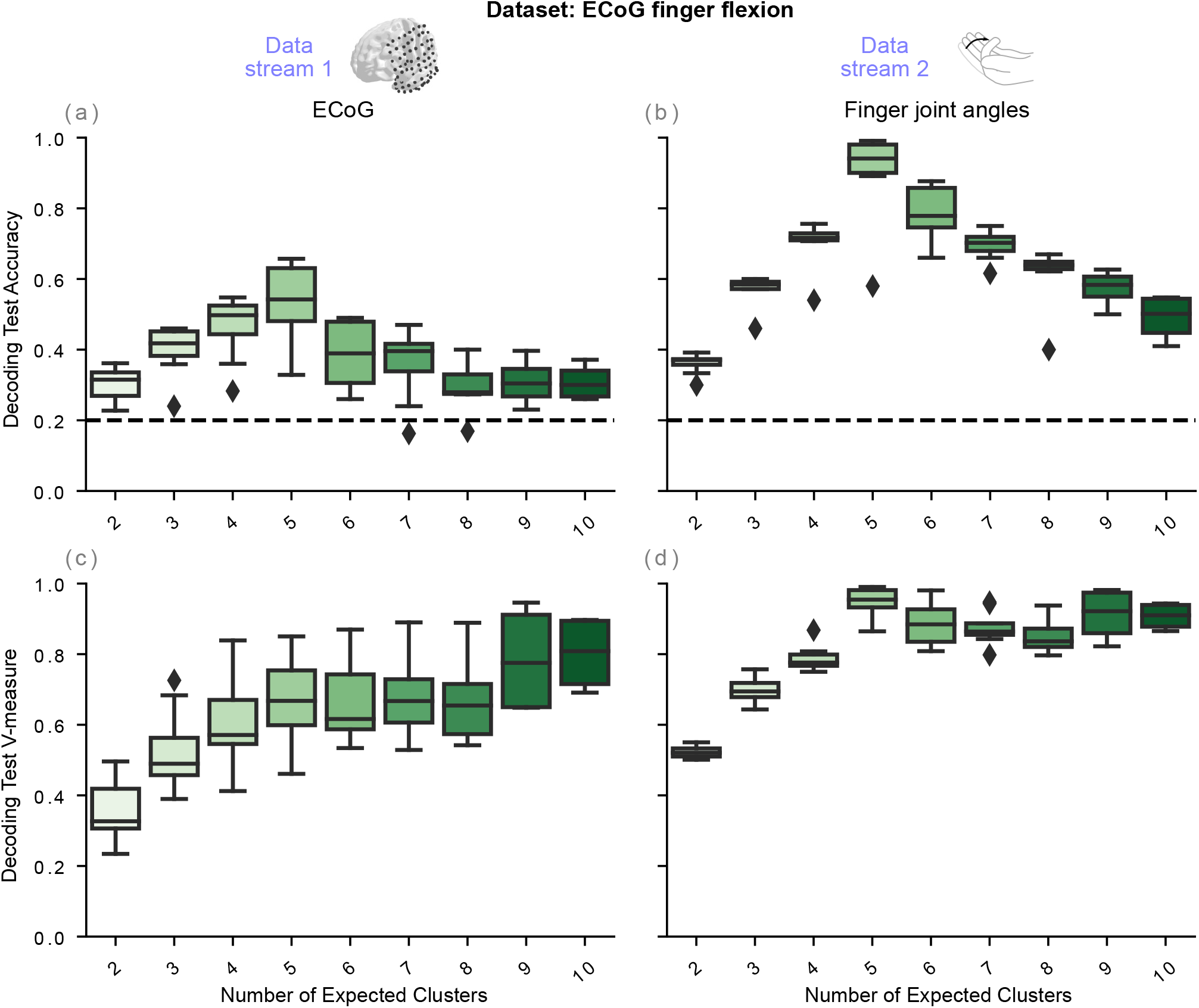
Effect of cluster number on cross-modal decoding performance. **(a–b)** Cross-modal test accuracy for the ECoG finger flexion dataset is highest when the number of clusters used to create the pseudo-labels matches the true number of classes (5). **(c–d)** In contrast, v-measure performance remains the same or slightly increases when the number of expected clusters is higher than the true number of classes. These findings demonstrate that the expected number of clusters should be carefully selected, especially if generalized decoding performance is desired. Test accuracy denotes average performance on withheld data over 10 folds, with random chance at 20%.

**Table S1:**
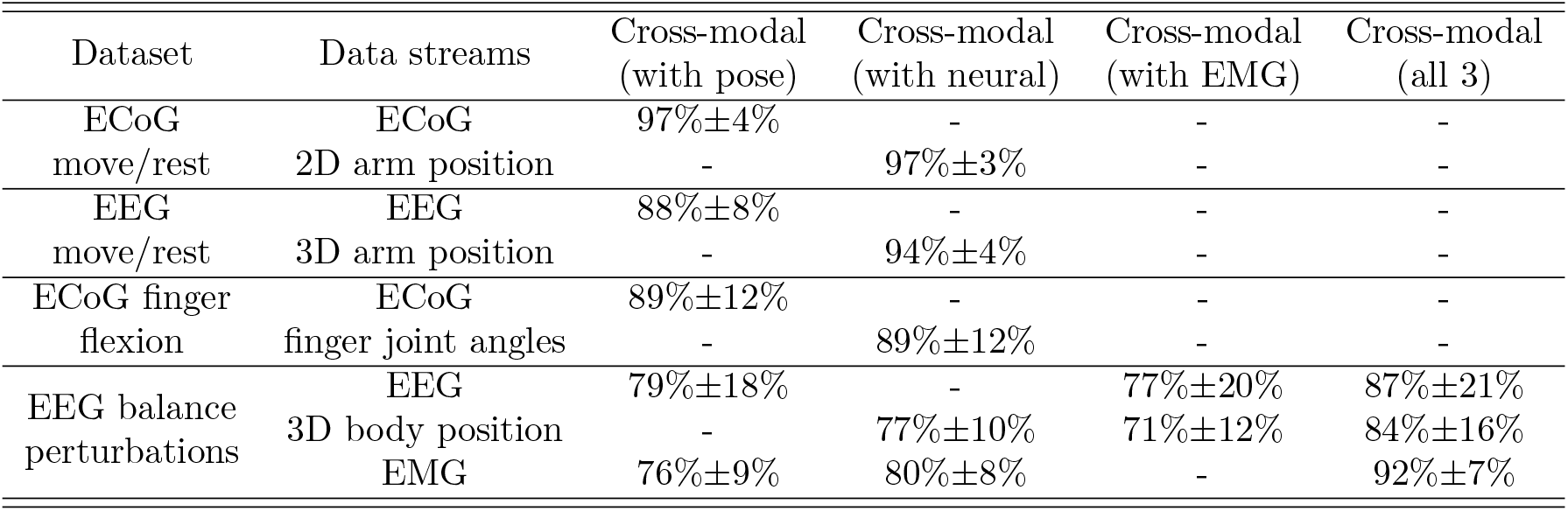
Decoding accuracy on the training data for cross-modal decoding. In most cases, we find that train accuracies (mean*±*SD) are quite similar across decoders that share pseudo-labels. These similarities suggest that much of the differences in cross-modal test accuracies among data streams is due to each decoder’s ability to generalize to unseen data.

**Table S2:**
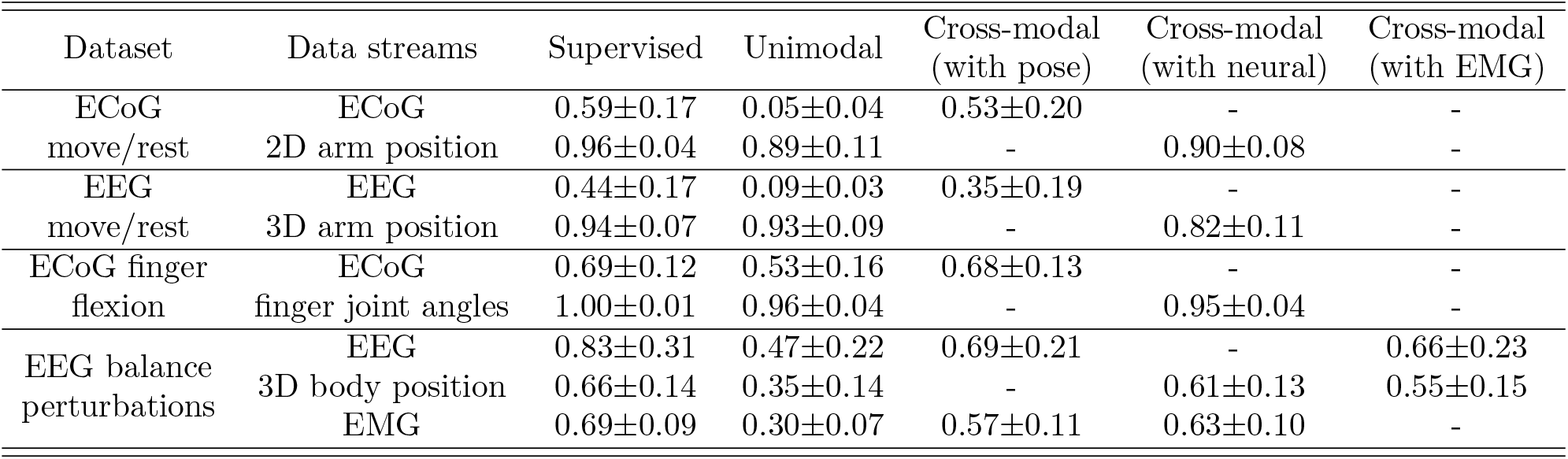
V-measure clustering performance for pairwise cross-modal decoding. V-measure assessment of cluster performance is shown for each dataset across participants (mean*±*SD). For nearly every data stream, cross-modal decoding has higher clustering performance than unimodal decoders, similar to decoding test accuracy.

**Table S3:** V-measure clustering performance for cross-modal decoding of three data streams. Similar to test accuracy, cluster performance across participants (mean*±*SD) for cross-modal models approaches or slightly exceeds supervised cluster performance.

| Dataset | Data streams | Supervised | Unimodal | Cross-modal<br>(all 3) |
| --- | --- | --- | --- | --- |
| EEG balance<br>perturbations | EEG | 0.83±0.31 | 0.47±0.22 | 0.80±0.26 |
| | 3D body position | 0.66±0.14 | 0.35±0.14 | 0.67±0.20 |
| | EMG | 0.69±0.09 | 0.30±0.07 | 0.72±0.11 |

**Table S4:** Decoding model hyperparameters. Decoding model hyperparameter values are shown for each dataset and data stream. We increased the number of spatial filters for datasets with more than 2 classes while reducing the number of temporal filters to avoid large differences in model size between datasets. SincNet filters and spectral power computation were either used (Y) or not used (N) for each data stream. Note that hyperparameter values were fixed across all self-supervised and supervised model types.

| Dataset | Data streams | Temporal filters | Spatial filters | Use SincNet filters | Compute spectral power |
| --- | --- | --- | --- | --- | --- |
| ECoG move/rest | ECoG | 19 | 2 | Y | Y |
|  | 2D arm position | 19 | 2 | Y | Y |
| EEG move/rest | EEG | 19 | 2 | Y | Y |
|  | 3D arm position | 19 | 2 | Y | Y |
| ECoG finger flexion | ECoG | 6 | 5 | Y | Y |
|  | finger joint angles | 6 | 5 | N | N |
| EEG perturbations | EEG | 6 | 3 | Y | Y |
|  | 3D body position | 6 | 3 | Y | Y |
|  | EMG | 6 | 3 | Y | Y |

## References

[1] Patrick D. Ganzer, Samuel C. Colachis, Michael A. Schwemmer, David A. Friedenberg, Collin F. Dunlap, Carly E. Swiftney, Adam F. Jacobowitz, Doug J. Weber, Marcia A. Bockbrader, and Gaurav Sharma. Restoring the sense of touch using a sensorimotor demultiplexing neural interface. Cell, 181(4):763–773.e12, 2020.

[2] Kai J Miller, Dora Hermes, and Nathan P Staff. The current state of electrocorticography-based brain–computer interfaces. Neurosurgical Focus, 49(1):E2, 2020.

[3] Ksenia Volkova, Mikhail A Lebedev, Alexander Kaplan, and Alexei Ossadtchi. Decoding movement from electrocorticographic activity: A review. Frontiers in neuroinformatics, 13, 2019.

[4] Soroush Niketeghad and Nader Pouratian. Brain machine interfaces for vision restoration: The current state of cortical visual prosthetics. Neurotherapeutics, 16(1):134–143, 2019.

[5] Stephanie Martin, Josédel R. Millán, Robert T. Knight, and Brian N. Pasley. The use of intracranial recordings to decode human language: Challenges and opportunities. Brain and Language, 193:73–83, 2019.

[6] Omid G Sani, Yuxiao Yang, Morgan B Lee, Heather E Dawes, Edward F Chang, and Maryam M Shanechi. Mood variations decoded from multi-site intracranial human brain activity. Nature Biotechnology, 36(10):954–961, 2018.

[7] David A Moses, Sean L Metzger, Jessie R Liu, Gopala K Anumanchipalli, Joseph G Makin, Pengfei F Sun, Josh Chartier, Maximilian E Dougherty, Patricia M Liu, Gary M Abrams, et al. Neuroprosthesis for decoding speech in a paralyzed person with anarthria. New England Journal of Medicine, 385(3):217–227, 2021.

[8] Alan D Degenhart, William E Bishop, Emily R Oby, Elizabeth C Tyler-Kabara, Steven M Chase, Aaron P Batista, and M Yu Byron. Stabilization of a brain–computer interface via the alignment of low-dimensional spaces of neural activity. Nature Biomedical Engineering, pages 1–14, 2020.

[9] Emily R Oby, Jay A Hennig, Aaron P Batista, M Yu Byron, and Steven M Chase. Intracortical brain–machine interfaces. In Neural Engineering, pages 185–221. Springer, 2020.

[10] Jennifer L Collinger, Robert A Gaunt, and Andrew B Schwartz. Progress towards restoring upper limb movement and sensation through intracortical brain-computer interfaces. Current Opinion in Biomedical Engineering, 8:84–92, 2018.

[11] Xiaotong Gu, Zehong Cao, Alireza Jolfaei, Peng Xu, Dongrui Wu, Tzyy-Ping Jung, and Chin-Teng Lin. Eeg-based brain-computer interfaces (bcis): A survey of recent studies on signal sensing technologies and computational intelligence approaches and their applications. arXiv preprint 2001.11337, 2020.

[12] Rajesh P. N. Rao. Brain-Computer Interfacing: An Introduction. Cambridge University Press, Cambridge, 2013.

[13] Chethan Pandarinath, K Cora Ames, Abigail A Russo, Ali Farshchian, Lee E Miller, Eva L Dyer, and Jonathan C Kao. Latent factors and dynamics in motor cortex and their application to brain–machine interfaces. Journal of Neuroscience, 38(44):9390–9401, 2018.

[14] Joshua I Glaser, Ari S Benjamin, Raeed H Chowdhury, Matthew G Perich, Lee E Miller, and Konrad P Kording. Machine learning for neural decoding. Eneuro, 7(4), 2020.

[15] Nima Bigdely-Shamlo, Jeremy Cockfield, Scott Makeig, Thomas Rognon, Chris La Valle, Makoto Miyakoshi, and Kay A Robbins. Hierarchical event descriptors (hed): semi-structured tagging for real-world events in large-scale eeg. Frontiers in neuroinformatics, 10:42, 2016.

[16] Pierre Karashchuk, Katie L Rupp, Evyn S Dickinson, Elischa Sanders, Eiman Azim, Bingni W Brunton, and John C Tuthill. Anipose: a toolkit for robust markerless 3d pose estimation. BioRxiv, 2020.

[17] Dongrui Wu, Yifan Xu, and Bao-Liang Lu. Transfer learning for eeg-based brain-computer interfaces: A review of progress made since 2016, 2020.

[18] Jan Van Erp, Fabien Lotte, and Michael Tangermann. Brain-computer interfaces: beyond medical applications. Computer, 45(4):26–34, 2012.

[19] Gan Huang, Guangquan Liu, Jianjun Meng, Dingguo Zhang, and Xiangyang Zhu. Model based generalization analysis of common spatial pattern in brain computer interfaces. Cognitive neurodynamics, 4(3):217–223, 2010.

[20] Mike X Cohen. Analyzing Neural Time Series Data: Theory and Practice, jan 2014.

[21] Longlong Jing and Yingli Tian. Self-supervised visual feature learning with deep neural networks: A survey. IEEE transactions on pattern analysis and machine intelligence, 2020.

[22] Xiao Liu, Fanjin Zhang, Zhenyu Hou, Li Mian, Zhaoyu Wang, Jing Zhang, and Jie Tang. Self-supervised learning: Generative or contrastive. IEEE Transactions on Knowledge and Data Engineering, 2021.

[23] Karine Lacourse, Ben Yetton, Sara Mednick, and Simon C Warby. Massive online data annotation, crowdsourcing to generate high quality sleep spindle annotations from eeg data. Scientific data, 7(1):1–14, 2020.

[24] Nancy Wang, Ali Farhadi, Rajesh Rao, and Bingni Brunton. Ajile movement prediction: Multimodal deep learning for natural human neural recordings and video. In Proceedings of the AAAI Conference on Artificial Intelligence, volume 32, 2018.

[25] Steven M Peterson, Zoe Steine-Hanson, Nathan Davis, Rajesh PN Rao, and Bingni W Brunton. Generalized neural decoders for transfer learning across participants and recording modalities. Journal of Neural Engineering, 18(2):026014, 2021.

[26] Humam Alwassel, Dhruv Mahajan, Bruno Korbar, Lorenzo Torresani, Bernard Ghanem, and Du Tran. Self-supervised learning by cross-modal audio-video clustering. Advances in Neural Information Processing Systems, 33, 2020.

[27] Tomas Mikolov, Ilya Sutskever, Kai Chen, Greg S Corrado, and Jeff Dean. Distributed representations of words and phrases and their compositionality. In Advances in neural information processing systems, pages 3111–3119, 2013.

[28] Jacob Devlin, Ming-Wei Chang, Kenton Lee, and Kristina Toutanova. Bert: Pre-training of deep bidirectional transformers for language understanding. arXiv preprint 1810.04805, 2018.

[29] Tom B Brown, Benjamin Mann, Nick Ryder, Melanie Subbiah, Jared Kaplan, Prafulla Dhariwal, Arvind Neelakantan, Pranav Shyam, Girish Sastry, Amanda Askell, et al. Language models are fewshot learners. arXiv preprint 2005.14165, 2020.

[30] Diederik P Kingma and Max Welling. Autoencoding variational bayes. arXiv preprint 1312.6114, 2013.

[31] Ian Goodfellow, Jean Pouget-Abadie, Mehdi Mirza, Bing Xu, David Warde-Farley, Sherjil Ozair, Aaron Courville, and Yoshua Bengio. Generative adversarial nets. Advances in neural information processing systems, 27, 2014.

[32] Kaiming He, Haoqi Fan, Yuxin Wu, Saining Xie, and Ross Girshick. Momentum contrast for unsupervised visual representation learning. In Proceedings of the IEEE/CVF Conference on Computer Vision and Pattern Recognition, pages 9729–9738, 2020.

[33] Ting Chen, Simon Kornblith, Mohammad Norouzi, and Geoffrey Hinton. A simple framework for contrastive learning of visual representations. In International conference on machine learning, pages 1597–1607. PMLR, 2020.

[34] Davide Chicco. Siamese neural networks: An overview. Artificial Neural Networks, pages 73–94, 2021.

[35] Hubert Banville, Omar Chehab, Aapo Hyvärinen, Denis-Alexander Engemann, and Alexandre Gramfort. Uncovering the structure of clinical eeg signals with self-supervised learning. Journal of Neural Engineering, 18(4):046020, 2021.

[36] Priya Goyal, Mathilde Caron, Benjamin Lefaudeux, Min Xu, Pengchao Wang, Vivek Pai, Mannat Singh, Vitaliy Liptchinsky, Ishan Misra, Armand Joulin, et al. Self-supervised pretraining of visual features in the wild. arXiv preprint 2103.01988, 2021.

[37] Jean-Bastien Grill, Florian Strub, Florent Altché, Corentin Tallec, Pierre H Richemond, Elena Buchatskaya, Carl Doersch, Bernardo Avila Pires, Zhaohan Daniel Guo, Mohammad Gheshlaghi Azar, et al. Bootstrap your own latent: A new approach to self-supervised learning. arXiv preprint 2006.07733, 2020.

[38] Mathilde Caron, Piotr Bojanowski, Armand Joulin, and Matthijs Douze. Deep clustering for unsupervised learning of visual features. In Proceedings of the European Conference on Computer Vision (ECCV), pages 132–149, 2018.

[39] Mathilde Caron, Ishan Misra, Julien Mairal, Priya Goyal, Piotr Bojanowski, and Armand Joulin. Unsupervised learning of visual features by contrasting cluster assignments. arXiv preprint 2006.09882, 2020.

[40] Kai J Miller. A library of human electrocorticographic data and analyses. Nature human behaviour, 3(11):1225–1235, 2019.

[41] Josef Parvizi and Sabine Kastner. Promises and limitations of human intracranial electroencephalography. Nature Neuroscience, 21(4):474–483, 2018.

[42] Kana Takaura, Naotsugu Tsuchiya, and Naotaka Fujii. Frequency-dependent spatiotemporal profiles of visual responses recorded with subdural ecog electrodes in awake monkeys: Differences between high- and low-frequency activity. NeuroImage, 124:557 – 572, 2016.

[43] Aysegul Gunduz, Peter Brunner, Amy Daitch, Eric C Leuthardt, Anthony L Ritaccio, Bijan Pesaran, and Gerwin Schalk. Neural correlates of visual-spatial attention in electrocorticographic signals in humans. Frontiers in human neuroscience, 5:89–89, 09 2011.

[44] Tobias Pistohl, Tonio Ball, Andreas Schulze-Bonhage, Ad Aertsen, and Carsten Mehring. Prediction of arm movement trajectories from ecog-recordings in humans. Journal of Neuroscience Methods, 167(1):105–114, 2008.

[45] S Raghu, Natarajan Sriraam, Erik D Gommer, Danny MW Hilkman, Yasin Temel, Shyam Vasudeva Rao, Alangar Satyaranjandas Hegde, and Pieter L Kubben. Cross-database evaluation of eeg based epileptic seizures detection driven by adaptive median feature baseline correction. Clinical Neurophysiology, 131(7):1567–1578, 2020.

[46] Alexander Ya Kaplan, Andrew A Fingelkurts, Alexander A Fingelkurts, Sergei V Borisov, and Boris S Darkhovsky. Nonstationary nature of the brain activity as revealed by eeg/meg: methodological, practical and conceptual challenges. Signal processing, 85(11):2190–2212, 2005.

[47] Scott Cole and Bradley Voytek. Cycle-by-cycle analysis of neural oscillations. Journal of neurophysiology, 122(2):849–861, 2019.

[48] Thomas Donoghue, Matar Haller, Erik J. Peterson, Paroma Varma, Priyadarshini Sebastian, Richard Gao, Torben Noto, Antonio H. Lara, Joni D. Wallis, Robert T. Knight, Avgusta Shestyuk, and Bradley Voytek. Parameterizing neural power spectra into periodic and aperiodic components. Nature Neuroscience, 23(12):1655–1665, 2020.

[49] Bradley Voytek, Mark A Kramer, John Case, Kyle Q Lepage, Zechari R Tempesta, Robert T Knight, and Adam Gazzaley. Age-related changes in 1/f neural electrophysiological noise. Journal of Neuroscience, 35(38):13257–13265, 2015.

[50] Agrita Dubey and Supratim Ray. Cortical electrocorticogram (ecog) is a local signal. Journal of Neuroscience, 39(22):4299–4311, 2019.

[51] Lau Troy M, Gwin Joseph T, and Ferris Daniel P. How many electrodes are really needed for eeg-based mobile brain imaging? Journal of Behavioral and Brain Science, 2012, 2012.

[52] Joseph Y Cheng, Hanlin Goh, Kaan Dogrusoz, Oncel Tuzel, and Erdrin Azemi. Subject-aware contrastive learning for biosignals. arXiv preprint 2007.04871, 2020.

[53] Demetres Kostas, Stephane Aroca-Ouellette, and Frank Rudzicz. Bendr: using transformers and a contrastive self-supervised learning task to learn from massive amounts of eeg data. arXiv preprint 2101.12037, 2021.

[54] Mostafa Neo Mohsenvand, Mohammad Rasool Izadi, and Pattie Maes. Contrastive representation learning for electroencephalogram classification. In Machine Learning for Health, pages 238–253. PMLR, 2020.

[55] Jinpei Han, Xiao Gu, and Benny Lo. Semi-supervised contrastive learning for generalizable motor imagery eeg classification. In 2021 IEEE 17th International Conference on Wearable and Implantable Body Sensor Networks (BSN), pages 1–4. IEEE, 2021.

[56] Steven M Peterson, Satpreet H Singh, Nancy XR Wang, Rajesh PN Rao, and Bingni W Brunton. Behavioral and neural variability of naturalistic arm movements. Eneuro, 2021.

[57] Roger Ratcliff, Marios G Philiastides, and Paul Sajda. Quality of evidence for perceptual decision making is indexed by trial-to-trial variability of the eeg. Proceedings of the National Academy of Sciences, 106(16):6539–6544, 2009.

[58] Colorado J Reed, Sean Metzger, Aravind Srinivas, Trevor Darrell, and Kurt Keutzer. Self-augment: Automatic augmentation policies for self-supervised learning. In Proceedings of the IEEE/CVF Conference on Computer Vision and Pattern Recognition, pages 2674–2683, 2021.

[59] Nikita Araslanov and Stefan Roth. Self-supervised augmentation consistency for adapting semantic segmentation. In Proceedings of the IEEE/CVF Conference on Computer Vision and Pattern Recognition, pages 15384–15394, 2021.

[60] Elnaz Lashgari, Dehua Liang, and Uri Maoz. Data augmentation for deep-learning-based electroencephalography. Journal of Neuroscience Methods, page 108885, 2020.

[61] Patrick Ofner, Andreas Schwarz, Joana Pereira, and Gernot R. Müller-Putz. Upper limb movements can be decoded from the time-domain of low-frequency eeg. PLOS ONE, 12(8):1–24, 08 2017.

[62] Kai J Miller, Dora Hermes, Christopher J Honey, Adam O Hebb, Nick F Ramsey, Robert T Knight, Jeffrey G Ojemann, and Eberhard E Fetz. Human motor cortical activity is selectively phase-entrained on underlying rhythms. PLoS Computational Biology, 8(9), 2012.

[63] Steven M Peterson and Daniel P Ferris. Differentiation in theta and beta electrocortical activity between visual and physical perturbations to walking and standing balance. eneuro, 5(4), 2018.

[64] Fiorenzo Artoni, Chiara Fanciullacci, Federica Bertolucci, Alessandro Panarese, Scott Makeig, Silvestro Micera, and Carmelo Chisari. Unidirectional brain to muscle connectivity reveals motor cortex control of leg muscles during stereotyped walking. Neuroimage, 159:403–416, 2017.

[65] Jinbiao Liu, Yixuan Sheng, and Honghai Liu. Corticomuscular coherence and its applications: a review. Frontiers in human neuroscience, 13:100, 2019.

[66] Evelyn Jungnickel, Lukas Gehrke, Marius Klug, Klaus Gramann, Hasan Ayaz, and Frédéric Dehais. Chapter 10 - MoBI—Mobile Brain/Body Imaging, pages 59–63. Academic Press, 2019.

[67] Yongtian He, Trieu Phat Luu, Kevin Nathan, Sho Nakagome, and Jose L Contreras-Vidal. A mobile brain-body imaging dataset recorded during treadmill walking with a brain-computer interface. Scientific data, 5(1):1–10, 2018.

[68] Alessandro Presacco, Ronald Goodman, Larry Forrester, and Jose Luis Contreras-Vidal. Neural decoding of treadmill walking from noninvasive electroencephalographic signals. Journal of neurophysiology, 106(4):1875–1887, 2011.

[69] Daniel G Wakeman and Richard N Henson. A multi-subject, multi-modal human neuroimaging dataset. Scientific data, 2(1):1–10, 2015.

[70] Grant Hanada. Mobile Brain and Body Imaging during Walking Motor Tasks. PhD thesis, University of Michigan, 2018.

[71] Nora Hollenstein, Cedric Renggli, Benjamin Glaus, Maria Barrett, Marius Troendle, Nicolas Langer, and Ce Zhang. Decoding eeg brain activity for multi-modal natural language processing. arXiv preprint 2102.08655, 2021.

[72] Raffaele Gravina, Parastoo Alinia, Hassan Ghasemzadeh, and Giancarlo Fortino. Multisensor fusion in body sensor networks: State-of-the-art and research challenges. Information Fusion, 35:68–80, 2017.

[73] Victor Javier Kartsch, Simone Benatti, Pasquale Davide Schiavone, Davide Rossi, and Luca Benini. A sensor fusion approach for drowsiness detection in wearable ultra-low-power systems. Information Fusion, 43:66–76, 2018.

[74] Yuki Markus Asano, Christian Rupprecht, and Andrea Vedaldi. Self-labelling via simultaneous clustering and representation learning. arXiv preprint 1911.05371, 2019.

[75] Marco Cuturi. Sinkhorn distances: Lightspeed computation of optimal transport. Advances in neural information processing systems, 26:2292–2300, 2013.

[76] Satpreet H Singh, Steven M Peterson, Rajesh PN Rao, and Bingni W Brunton. Mining naturalistic human behaviors in long-term video and neural recordings. Journal of Neuroscience Methods, 358:109199, 2021.

[77] Vernon J Lawhern, Amelia J Solon, Nicholas R Waytowich, Stephen M Gordon, Chou P Hung, and Brent J Lance. Eegnet: a compact convolutional neural network for eeg-based brain–computer interfaces. Journal of neural engineering, 15(5):056013, 2018.

[78] Andrew Rosenberg and Julia Hirschberg. V-measure: A conditional entropy-based external cluster evaluation measure. In Proceedings of the 2007 joint conference on empirical methods in natural language processing and computational natural language learning (EMNLP-CoNLL), pages 410–420, 2007.

[79] Kai Han, Andrea Vedaldi, and Andrew Zisserman. Learning to discover novel visual categories via deep transfer clustering. In Proceedings of the IEEE/CVF International Conference on Computer Vision, pages 8401–8409, 2019.

[80] Milton Friedman. The use of ranks to avoid the assumption of normality implicit in the analysis of variance. Journal of the american statistical association, 32(200):675–701, 1937.

[81] William Jay Conover. Practical nonparametric statistics, volume 350. John Wiley & Sons, 1998.

[82] Yoav Benjamini and Yosef Hochberg. Controlling the false discovery rate: a practical and powerful approach to multiple testing. Journal of the Royal statistical society: series B (Methodological), 57(1):289–300, 1995.

[83] Y Tran, RA Thuraisingham, N Wijesuriya, HT Nguyen, and A Craig. Detecting neural changes during stress and fatigue effectively: a comparison of spectral analysis and sample entropy. In 2007 3rd International IEEE/EMBS Conference on Neural Engineering, pages 350–353. IEEE, 2007.

[84] Laura B Baucom, Douglas H Wedell, Jing Wang, David N Blitzer, and Svetlana V Shinkareva. Decoding the neural representation of affective states. Neuroimage, 59(1):718–727, 2012.

[85] Jordan Muraskin, Truman R Brown, Jennifer M Walz, Tao Tu, Bryan Conroy, Robin I Goldman, and Paul Sajda. A multimodal encoding model applied to imaging decision-related neural cascades in the human brain. Neuroimage, 180:211–222, 2018.

[86] Sarwat Fatima and Awais M Kamboh. Decoding brain cognitive activity across subjects using multimodal m/eeg neuroimaging. In 2017 39th Annual International Conference of the IEEE Engineering in Medicine and Biology Society (EMBC), pages 3224–3227. IEEE, 2017.

[87] Federico Gennaro and Eling D De Bruin. Assessing brain–muscle connectivity in human locomotion through mobile brain/body imaging: Opportunities, pitfalls, and future directions. Frontiers in public health, 6:39, 2018.

[88] Steven M Peterson, Emily Furuichi, and Daniel P Ferris. Effects of virtual reality high heights exposure during beam-walking on physiological stress and cognitive loading. PloS one, 13(7):e0200306, 2018.

[89] Kaiming He, Xiangyu Zhang, Shaoqing Ren, and Jian Sun. Deep residual learning for image recognition. In Proceedings of the IEEE conference on computer vision and pattern recognition, pages 770–778, 2016.

[90] Sepp Hochreiter and Jürgen Schmidhuber. Long short-term memory. Neural computation, 9(8):1735–1780, 1997.

[91] Kai-Nan An. Kinematic analysis of human movement. Annals of biomedical engineering, 12(6):585–597, 1984.

[92] Wei Fu and Patrick O Perry. Estimating the number of clusters using cross-validation. Journal of Computational and Graphical Statistics, 29(1):162–173, 2020.

[93] Luca Randazzo, Iñaki Iturrate, Ricardo Chavarriaga, Robert Leeb, and Josédel R Millán. Detecting intention to grasp during reaching movements from eeg. In 2015 37th Annual International Conference of the IEEE Engineering in Medicine and Biology Society (EMBC), pages 1115–1118. IEEE, 2015.

[94] Rami Alazrai, Hisham Alwanni, and Mohammad I Daoud. Eeg-based bci system for decoding finger movements within the same hand. Neuroscience letters, 698:113–120, 2019.

[95] Morten L Kringelbach and Gustavo Deco. Brain states and transitions: insights from computational neuroscience. Cell Reports, 32(10):108128, 2020.

[96] Talita Peixoto Pinto, Maitê Mello Russo Ramos, Thiago Lemos, Claudia Domingues Vargas, and Luis Aureliano Imbiriba. Is heart rate variability affected by distinct motor imagery strategies? Physiology & behavior, 177:189–195, 2017.

[97] Elke Heremans, Werner F Helsen, and Peter Feys. The eyes as a mirror of our thoughts: quantification of motor imagery of goal-directed movements through eye movement registration. Behavioural Brain Research, 187(2):351–360, 2008.

[98] F Lebon, David Rouffet, C Collet, and A Guillot. Modulation of emg power spectrum frequency during motor imagery. Neuroscience letters, 435(3):181–185, 2008.

[99] Omid G Sani, Yuxiao Yang, Morgan B Lee, Heather E Dawes, Edward F Chang, and Maryam M Shanechi. Mood variations decoded from multi-site intracranial human brain activity. Nature biotechnology, 36(10):954–961, 2018.

[100] Sharlene N Flesher, John E Downey, Jeffrey M Weiss, Christopher L Hughes, Angelica J Herrera, Elizabeth C Tyler-Kabara, Michael L Boninger, Jennifer L Collinger, and Robert A Gaunt. A brain-computer interface that evokes tactile sensations improves robotic arm control. Science, 372(6544):831–836, 2021.

[101] Mirco Ravanelli and Yoshua Bengio. Speaker recognition from raw waveform with sincnet. In 2018 IEEE Spoken Language Technology Workshop (SLT), pages 1021–1028. IEEE, 2018.

[102] Steven M Peterson, Satpreet H Singh, Benjamin Dichter, Michael Scheid, Rajesh PN Rao, and Bingni W Brunton. Ajile12: Long-term naturalistic human intracranial neural recordings and pose. bioRxiv, 2021.

[103] Inderjeet Mani and I Zhang. knn approach to unbalanced data distributions: a case study involving information extraction. In Proceedings of workshop on learning from imbalanced datasets, volume 126. ICML United States, 2003.

